## Supplementary Material for "Intracellular competitions reveal the combination of active partitioning and addiction mechanisms as most advantageous for low-copy plasmid fitness"

Effe et al.

**This PDF file includes:**
Supplementary text: information on the mathematical model of plasmid segregation
Figures S1 to S4
List of tables S1 to S5
List of datasets S1 to S4

### Supplemental text: model description

We simulate the persistence dynamics and the head-to-head competitions of a multicopy plasmid with various genetic variants using a random segregation model[1,2], which we have extended for plasmid stability mechanisms. In the extended model, bacterial cells initially carry a maximum of two plasmid variants. The diagram in Figure 3A shows the life cycle for a heteroplasmic cell with two plasmid copies one encoding an active partition system (P) and the other one a toxin-antitoxin system (T), respectively.

The life cycle includes four steps: 1) plasmid replication, 2) monomerization, 3) assortment to daughter cells, and 4) post-segregational killing. Plasmid copies replicate following a Pólya urn model, where copies are randomly selected and their replicates are returned to the urn (cell) before the next plasmid replication. The plasmid copy number, $n_c$, is jointly controlled for all plasmid variants. Specifically, plasmid replication ceases when a cell contains $2n_c$ plasmid copies, regardless of the number of plasmid copies at cell birth.

In the second step, unsuccessful monomerization of plasmid replicates due incomplete plasmid separation (clustering) or plasmid fusion can hinder faithful inheritance of all plasmid replicates[3]. We assume that a new plasmid replicate is eliminated with a probability $p_{\rm def}$, as replicates that are part of a plasmid multimer or cluster are not subject to random segregation. We do not consider plasmid fusion of genetically different plasmid variants.

In the third step, plasmid copies are randomly assorted to one daughter cell during cell division, except for plasmids encoding an active partition system (P). Copies of the P (and PT) plasmid variant form pairs and the plasmid copies of each pair are assorted to separate daughter cells with a probability $p_{\rm par}$. Otherwise, with a probability $1-p_{\rm par}$, the plasmids are randomly distributed to the daughter cells.

Finally, in the fourth step, the plasmid-encoded toxin-antitoxin system can cause cell death of newly created daughter cells that lack the antitoxin-encoding plasmid (T). Daughter cells without the T-plasmid are eliminated with a probability $p_{\rm eff}$, which represents the effectiveness of post-segregational killing by the toxin-antitoxin system. Conversely, with a probability $1-p_{\rm eff}$, a daughter cell lacking the antitoxin-encoding T-plasmid survives and proliferates, producing new offspring.

#### Mathematical description

To make our model mathematically tractable, we denote a cell's genotype (cell-type) by a tuple $(i_{p_1}, i_{p_2})$, where $p_1, p_2 \in \{U, S, P, T, PT\}$, $p_1 \neq p_2$ denote the two different plasmid variants involved, and $i_{p_1}$ and $i_{p_2}$ describe the abundance of the respective plasmid variants. Note that $i_{p_1}$ and $i_{p_2}$ are the copy numbers at cell birth, which may change during a cell's life cycle. The cell-type $(i_{p_1}, i_{p_2}) = (1,1)$, where $p_1 = P$, $p_2 = T$, shown in the example of Figure 3A, describes a cell with one plasmid copy of the P-plasmid and one plasmid copy of the T-plasmid at cell birth. For cells carrying a plasmid with copy number $n_{\rm c}$, we have to consider $\frac{(2n_c+1)(2n_c+2)}{2}$ cell types, arising from the fact that there is one possible cell-type for the plasmid-free cell-type, two possible types for cells with one plasmid copy, three types for cells with two plasmid copies, and so on. The maximal number of plasmid replicates is $2n_c$ as the copy number is strictly controlled in our model, such that random plasmid assortment allows at maximum $2n_c$ copies to be distributed to one newly created daughter cell.

To analyze the plasmid evolution, we need to compute the probabilities, $p_{i\to j}$, that a cell of type $i = (i_{p_1}, i_{p_2})$ gives rise to a daughter cell of type $j = (j_{p_1}, j_{p_2})$. In the following, we only describe the scenario, where at most one of the two plasmid variants encodes a partioning system, i.e., we exclude $p_1 = P$, $p_2 = PT$ or vice-versa, as the latter is not of interest for our analyses since plasmids with the same stability determinant would anyway recombine frequently. In case one plasmid variant encodes a partitioning system (P or PT), we denote its plasmid copy number by $i_{p_1}$, while the second plasmid, $p_2$, lacks the partitioning system. Let $J_{p_1}^{\mathrm{rep}}$ and $J_{p_2}^{\mathrm{rep}}$ denote the random numbers that describe the number of replicates after plasmid replication,

$$J_{p_1}^{\mathrm{rep}} - i_{p_1} \sim \mathrm{BetaBin}(n = 2n_c - i_{p_1} - i_{p_2}, \alpha = i_{p_1}, \beta = i_{p_2}) \tag{1}$$

and $J_{p_2}^{\mathrm{rep}} = 2n_c - J_{p_1}^{\mathrm{rep}}$. $\mathrm{BetaBin}(n, \alpha, \beta)$ denotes a random number following a beta-binomial distribution, which has a probability mass function $\mathrm{PMF}(X = k) = \binom{n}{k}\frac{\mathrm{B}(k+\alpha, n-k+\beta)}{\mathrm{B}(\alpha,\beta)}$, where $B(m, n) = \frac{(m-1)!(n-1)!}{(m+n-1)!}$. For fixed $J_{p_1}^{\mathrm{rep}}$ and $J_{p_2}^{\mathrm{rep}}$, the numbers of plasmid replicates of variant 1 and 2 after monomerization, which we denote by $J_{p_1}^{\mathrm{mon}}$ and $J_{p_2}^{\mathrm{mon}}$, respectively, are binomially distributed:

$$J_{p_1(p_2)}^{\mathrm{mon}} | J_{p_1(p_2)}^{\mathrm{rep}} \sim \mathrm{Bin}(n = J_{p_1(p_2)}^{\mathrm{rep}} - i_{p_1(p_2)}, p = 1 - p_{\mathrm{def}}^{p_1(p_2)}) + i_{p_1(p_2)} \tag{2}$$

We only describe the genotype of one focal daughter cell, denoted by $j$, as we are interested only in the expected cell-type frequencies, which is based on the assumption that cell populations are large. Plasmid assortment may be influenced by active partitioning yet the number of plasmid replicates distributed to one daughter cell in absence of a partitioning system (as for variant 2) is binomially distributed, i.e.,

$$J_{p_2}^{\mathrm{div}} | J_{p_2}^{\mathrm{mon}} \sim \mathrm{Bin}(n = J_{p_2}^{\mathrm{mon}}, p = 1/2) \tag{3}$$

For plasmid variant 1 possibly encoding a partitioning system, we need to take into account the assortment of 1. the plasmid pairs ($\lfloor J_{p_1}^{\mathrm{mon}}/2 \rfloor$) and 2. the remaining single plasmid copy in case $J_{p_1}^{\mathrm{mon}}$ is an odd number: An elegant mathematical way to describe the probability distribution of $J_{p_1}^{\mathrm{div}}$ is by the use of probability generating functions, where

$$P(J_{p_1}^{\mathrm{div}} = k | J_{p_1}^{\mathrm{mon}}) = \frac{G^{(k)}(0; J_{p_1}^{\mathrm{mon}})}{k!} \tag{4}$$

with $G(z; J_{p_1}^{\mathrm{mon}}) = \left(\frac{1 - p_{\mathrm{sep},1}}{4} z^2 + \frac{p_{\mathrm{sep},1} + 1}{2} z + \frac{1 - p_{\mathrm{sep},1}}{4}\right)^{\lfloor J_{p_1}^{\mathrm{mon}}/2 \rfloor} \left(\frac{1}{2} z + \frac{1}{2}\right)^{\left(J_{p_1}^{\mathrm{mon}} - 2\lfloor J_{p_1}^{\mathrm{mon}}/2 \rfloor\right)}$ (5)

The polynomial coefficients in the first brackets describe the probability that two, one, or zero replicates of any pair are assorted to one daughter cell while the second factor accounts for the assortment of the possibly remaining single copy in case $J_{p_1}^{\mathrm{mon}}$ is odd. Note that, in case of $p_{\mathrm{sep},1} = 0$ (no active partitioning), we have $G^{(k)}(z) = (\frac{1}{2}z + \frac{1}{2})^{J_{p_1}^{\mathrm{mon}}}$, and thus $P(J_{p_1}^{\mathrm{div}} = k) = \binom{n}{k}(\frac{1}{2})^k(\frac{1}{2})^{n-k}$, such that $J_{p_1}^{\mathrm{div}}$ is binomially distributed like $J_{p_2}^{\mathrm{div}}$. The probability that the focal daughter cell has type $j = (J_{p_1}, J_{p_2})$ when a cell of type $i$ divides yields

$$\begin{aligned} p_{i \to j} = &\sum_{\substack{k_1 = i_{p_1}, \ldots, 2n_c - i_{p_2} \\ k_2 = i_{p_2}, \ldots, 2n_c - i_{p_1}}} P((J_{p_1}^{\mathrm{rep}}, J_{p_2}^{\mathrm{rep}}) = (k_1, k_2)) \\ &\sum_{\substack{l_1 = i_{p_1}, \ldots, k_1 \\ l_2 = i_{p_2}, \ldots, k_2}} P(J_1^{\mathrm{mon}} = l_1 | J_{p_1}^{\mathrm{rep}} = k_1) P(J_2^{\mathrm{mon}} = l_2 | J_{p_2}^{\mathrm{rep}} = k_2) \\ &P(J_1^{\mathrm{div}} = J_{p_1} | J_1^{\mathrm{mon}} = l_1) P(J_2^{\mathrm{div}} = J_{p_2} | J_2^{\mathrm{mon}} = l_2). \end{aligned} \tag{6}$$

In the next step, we compute the expected change of cells of type $j$ when an $i$-type cell divides. After cell division, new cells in our model are eliminated with a probability $p_{\mathrm{eff}}$ by the toxin-antitoxin system, in case they lack the antitoxin system encoded on the plasmid while their parental cell carried it. We have to distinguish two cases for which we calculate the probability of cell survival, $p_{i\to j}^{\mathrm{surv}}$, for a cell of type $j$ originating from an $i$-type cell: 1. Both plasmid variants code for the toxin-antitoxin system,

$$p_{i\to j}^{\mathrm{surv}} = \begin{cases} 1 - p_{\mathrm{eff}} & \text{if } (i_{p_1} > 0 \text{ or } i_{p_2} > 0) \text{ and } (J_{p_1} = 0 \text{ or } J_{p_2} = 0) \\ 1 & \text{otherwise;} \end{cases} \tag{7}$$

2. only plasmid variant 1 (or 2) codes for the toxin-antitoxin system,

$$p_{i\to j}^{\mathrm{surv}} = \begin{cases} 1 - p_{\mathrm{eff}} & \text{if } i_{1(2)} > 0 \text{ and } j_{1(2)} = 0 \\ 1 & \text{otherwise.} \end{cases} \tag{8}$$

We can compute the expected number of $j$-type cells that were produced and survived after cell division of an $i$-type cell as

$$2p_{i\to j}p_{i\to j}^{\mathrm{surv}} =: A_{ji}, \tag{9}$$

where $i$ and $j$ denote the multi-indices for the cell-types $(i_{p_1}, i_{p_2})$ and $(J_{p_1}, J_{p_2})$, respectively. Let us define the matrix

$$\mathbf{A} := (A_{ji})_{\tilde{j}\tilde{i}}, \tag{10}$$

where $\tilde{j}$, $\tilde{i}$ are indices for all the cell-types and $j = (j_{p_1}, j_{p_2}), i = (i_{p_1}, i_{p_2})$ are the multi-indices corresponding to these type indices, $\tilde{j}, \tilde{i}$, respectively. The expected number of cells after $g$ generations for a starting population $\mathbf{n}_0 = (n_{\tilde{i}})_{\tilde{i}}$, where the vector elements $n_{\tilde{i}}$ denote the initial number of cells of type $i$, $i_{p_1} \in \{0, \ldots, 2n_c\}, i_{p_2} \in \{0, 2n_c - i_{p_1}\}$, can be computed as

$$\mathbf{A}^g \mathbf{n}_0 = \underbrace{\mathbf{A} \cdot \mathbf{A} \ldots \mathbf{A}}_{g-\text{times}} \mathbf{n}_0, \tag{11}$$

and, for interpolation between two generations, i.e., at non-integer generation times $g$, we use the power series, $\mathbf{A}^g = \sum_{k=0}^{\infty} \frac{\mathrm{d}^k(x^g)}{\mathrm{d}x^k}\Big|_{x=1} (\mathbf{A} - \mathbf{I})^k$, where $\mathbf{I}$ denotes the identity matrix. In populations without a toxin-antitoxin encoding (TA) plasmids, all cells survive after cell division ($p_{i\to j}^{\mathrm{surv}} = 1$ for all $i, j$). Therefore, the population doubles each generation such that the proportions of cell-types after $g$ generations can be directly calculated by

$$\mathbf{x}(g) = \left(\frac{\mathbf{A}}{2}\right)^g \mathbf{x}(0), \tag{12}$$

where $\mathbf{x}(0) = \frac{\mathbf{n}_0}{|\mathbf{n}_0|}$. In populations with the TA plasmid, some proportion of the $2|\mathbf{n}_0|$ cells after cell division is subject to post-segregational killing, which we have to take into account when computing the proportions of cell types. To compute the expected proportions of cell types after $g$ generations, $\mathbf{x}(g)$, we divide the cell-type abundances by the expected number of cells,

$$\mathbf{x}(g) = \frac{\mathbf{A}^g \mathbf{n}_0}{|\mathbf{A}^g \mathbf{n}_0|} = \frac{\mathbf{A}^g \mathbf{x}_0}{|\mathbf{A}^g \mathbf{x}_0|} \tag{13}$$

For populations with only one genetic variant of the plasmid, the dimensionality of the matrix $\mathbf{A}$ can be reduced by omitting cell-types that describe the abundances of the second plasmid variant. We define the reduced matrix by

$$\mathbf{A}^{(1)} = (A_{(j_{p_1},0),(i_{p_1},0)})_{j_{p_1}, i_{p_1} \in \{0,\ldots,2n_c\}}, \tag{14}$$

where the matrix elements according to Eq. (9), (6), and (7) are

$$A_{(j_{p_1},0),(i_{p_1},0)} = 2p_{(i_{p_1},0)\to(j_{p_1},0)} p_{(i_{p_1},0)\to(j_{p_1},0)}^{(\mathrm{surv})} \tag{15}$$

with

$$p_{(i_{p_1},0)\to(j_{p_1},0)} = \sum_{l_1}^{2n_c} P(J_1^{\mathrm{mon}} = l_1 | J_{p_1}^{\mathrm{rep}} = 2n_c) P(J_1^{\mathrm{div}} = J_{p_1} | J_1^{\mathrm{mon}} = l_1), \tag{16}$$

$$p^{(\mathrm{surv})}_{(i_{p_1},0)\to(J_{p_1},0)} = \begin{cases} 1 - p_{\mathrm{eff}} & \text{if } i_{p_1} > 0 \text{ and } J_{p_1} = 0, \\ 1 & \text{otherwise.} \end{cases} \tag{17}$$

To compute the persistence of the plasmid in a population, we evaluate the cell-type abundances of cells carrying from zero up to $2n_c$ plasmid copies given by

$$\mathbf{n}^{(1)} = (n_0^{(1)}, n_1^{(1)}, \dots, n_{2n_c}^{(1)}) \tag{18}$$

and calculate there proportions, $\mathbf{x}^{(1)} = \frac{\mathbf{n}_0^{(1)}}{|\mathbf{n}_0^{(1)}|}$, at generation $g$, by

$$\mathbf{x}^{(1)}(g) = \frac{\mathbf{A}^{(1)g}\mathbf{x}_0^{(1)}}{|\mathbf{A}^{(1)g}\mathbf{x}_0^{(1)}|}. \tag{19}$$

or, in case of $p_{\mathrm{eff}} = 0$, (analogue to Eq. (12)) by

$$\mathbf{x}^{(1)}(g) = \left(\frac{\mathbf{A}^{(1)}}{2}\right)^g \mathbf{x}^{(1)}(0). \tag{20}$$

In the latter case, we can make use of the Eigenvalue decomposition

$$\mathbf{x}^{(1)}(g) = \left(\frac{\lambda_0}{2}\right)^g x_0\mathbf{v}_0 + \left(\frac{\lambda_1}{2}\right)^g x_1\mathbf{v}_1, \tag{21}$$

where $\lambda_0$, $\lambda_1$, $\lambda_2$, ..., $\lambda_{2n_c}$ denote the eigenvalues of $\mathbf{A}^{(1)}$ and $\mathbf{v}_0$, $\mathbf{v}_1$, $\mathbf{v}_2$, ..., $\mathbf{v}_{2n_c}$ are the corresponding eigenvectors. Note that $\lambda_2 = \ldots = \lambda_{2n_c} = 0$; therefore, the eigenvalue composition is a linear combination of $\mathbf{v}_0$ and $\mathbf{v}_1$. From the construction of the model and the resulting shape of $\mathbf{A}^{(1)}$, where $A_{i0}^{(1)} = 2$ if $i = 0$ and $A_{ij} = 0$ otherwise (i.e., in words, plasmid-free cells give rise to plasmid-free cells), we can assess $\lambda_0 = 2$ with the eigenvector $\mathbf{v}_1 = (1, 0, \dots)$. Moreover, there are $2n_c - 1$ eigenvalues equal to zero, $\lambda_2 = \lambda_3 = \cdots = \lambda_{2n_c} = 0$, as $A_{ij}^{(1)} = A_{ik}^{(1)}$ for any $i, k > 0$ which comes from the fact that the exact cell-type of plasmid-host cells ($i > 0$), which denotes the plasmid copies at cell birth, is irrelevant as the plasmid is replicated until $2n_c$ copies are reached. To compute the plasmid loss, we need to analyse $\lambda_1$ and correspoding the eigenvector $\mathbf{v}_1 = (v_{1,0}, v_{1,1}, \dots, v_{1,2n_c})$, which we normalize such that $v_{1,0} = -1$ such that $v_{1,1} + \cdots + v_{1,2n_c} = 1$. We can thus write any initial population that is consisting only of plasmid hosts, $\mathbf{x}^{(1)}(0)$, where $x_0^{(1)}(0) = 0$ as a superposition, with $x_0 = x_1 = 1$

$$\mathbf{x}^{(1)}(0) = \mathbf{v}_0 + \mathbf{v}_1 + x_2\mathbf{v}_2 + x_{2n_c}\mathbf{v}_{2n_c} \tag{22}$$

since the eigenvectors $\mathbf{v}_2$, ..., $\mathbf{v}_{2n_c}$ only change the composition of plasmid-host cell types ($\sum_j^{2n_c} v_{ij} = 0$ for all $i \geq 2$). This allows us to obtain the plasmid-host proportion for any generation $g > 0$ by

$$x_h(g) := \sum_{i=1}^{2n_c} x^{(1)}(g)_i = \sum_{i=1}^{2n_c} \left(\frac{\lambda_1}{2}\right)^g x_1\mathbf{v}_1 = \left(\frac{\lambda_1}{2}\right)^g. \tag{23}$$

The eigenvalue $\lambda_1 = \lambda_1(p_{\mathrm{def}}, p_{\mathrm{sep}})$ is a function of the parameters $p_{\mathrm{def}}$ and $p_{\mathrm{sep}}$, which can be calculated analytically (to our knowledge only) for $p_{\mathrm{def}} = p_{\mathrm{sep}} = 0$,

$$\lambda_1 = 2(1 - \frac{1}{2^{2n_c}}); \tag{24}$$

note that $r^{(S)} = 1 - \frac{1}{2^{2n_c}}$ is simply the segregation rate for completely randomly segregating plasmids. Thus, we compute $\lambda_2$ numerically if $(p_{\mathrm{def}} > 0$ or $p_{\mathrm{sep}} > 0.$

The plasmid-host proportion can also be directly calculate from Eq. (19), which we make use of for plasmids encoding the toxin-antitoxin system ($p_{\mathrm{eff}} > 0$), by

$$x_h(g) = \sum_{i=1}^{2n_c} x^{(1)}(g)_i. \tag{25}$$

### Estimation of parameters

For the estimation of the parameters, we fitted pCON$^{n}/_{c}$ persistence data using either $p_{\mathrm{def}}$, $p_{\mathrm{par}}$, or $p_{\mathrm{eff}}$ as a free parameter while leaving the other parameters fixed. We started using the pCON$^{n}/_{c}$-U persistence data to estimate $p_{\mathrm{def}}$ and constrained $p_{\mathrm{par}} = 0$ and $p_{\mathrm{eff}} = 0$. To assess the generation times corresponding to the persistence data acquired from serial transfer experiment, we computed the generations passed between two consecutive transfers, $d-1$ and $d$, from the end population size, $N_d$, and the start population size $N_{d-1}/b$ with $b$ denoting the bottleneck factor, which yields

$$\log_2\left(\frac{N_d}{N_{d-1}/b}\right) \tag{26}$$

The generation time at any day in the serial transfer experiment, $d = 1, \ldots$, can be cumulated by

$$g_d = g_{d-1} + \log_2\left(\frac{N_d}{N_{d-1}/b}\right). \tag{27}$$

Using the experimentally measured proportions, $x_{h,d}^{(\mathrm{exp})}(g_d)$, of cells carrying the pCON$^{n}/_{c}$-U plasmid (plasmid host cells), we calculated the best fitting parameter $p_{\mathrm{def}}^{(\mathrm{fit})}$ using the model function

$$x_h(g) = \sum_{i=1}^{2n_c} \left(\frac{\mathbf{A}^{(1)g}\hat{\mathbf{e}}_1}{|\mathbf{A}^{(1)g}\hat{\mathbf{e}}_1|}\right)_i, \tag{28}$$

where $\hat{\mathbf{e}}_1 = (0, 1, 0, \ldots)$ denotes the unit vector along the first dimension reflecting the initial condition of a homogeneous population of plasmid hosts (with one plasmid copy per cell). For the estimation of the other parameters, $p_{\mathrm{par}}$ and $p_{\mathrm{eff}}$, we constrained the model by $p_{\mathrm{def}} = p_{\mathrm{def}}^{(\mathrm{fit})}$ and then fitted the pCON$^{n}/_{c}$-P persistence data with $p_{\mathrm{par}}$ as a free parameter (while $p_{\mathrm{eff}} = 0$) and the pCON$^{n}/_{c}$-T persistence data with $p_{\mathrm{eff}}$ as a free parameter (while $p_{\mathrm{par}} = 0$).

### Plasmid segregation rate and half life

From the eigenvalue $\lambda_2$, we can also compute the segregation rate by

$$r = 2(1 - \lambda_2/2), \tag{29}$$

which describes the probability that plasmid segregation leads to the creation a plasmid-free cell as well the half life of a plasmid

$$\lambda^{(1/2)} = \frac{\log(1/2)}{\log(\lambda_2)} \tag{30}$$

For the toxin-antitoxin encoding plasmid variants, we compute the half life of a plasmid by numerically solving $x_h(g) = \frac{1}{2}$ using Eq. (28).

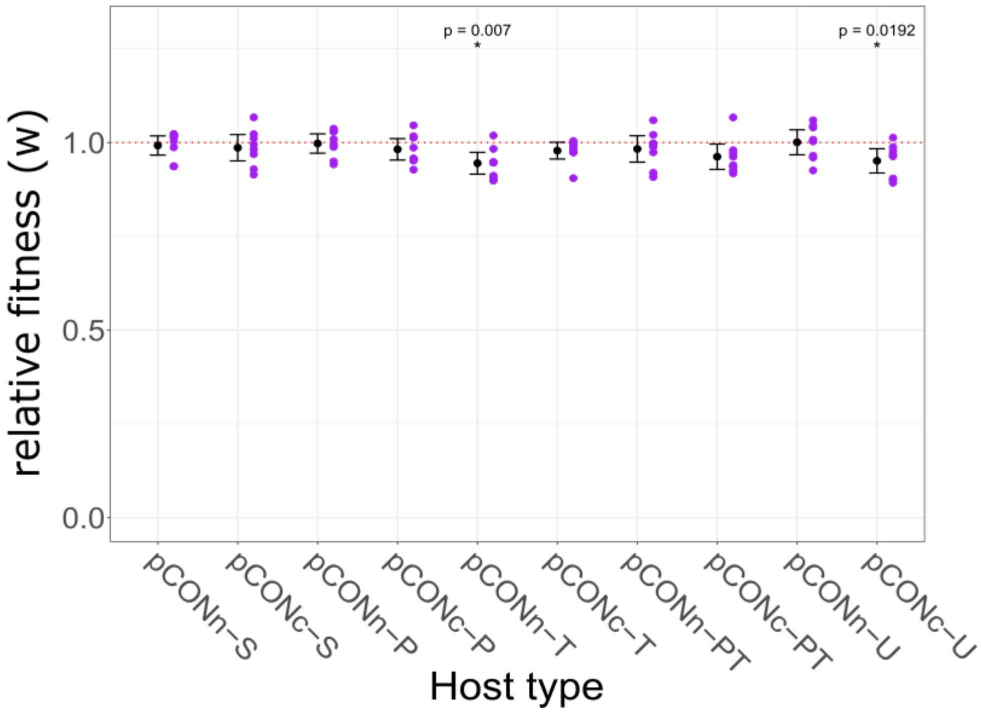


**Fig. S1. Fitness effect of the model plasmids on their host.**

Relative fitness of *E. coli* MG1655 *recA*::*tetA* hosting the model plasmids in comparison to an *E. coli* MG1655 *recA*::*tetA* TmR strain. Depicted here are the results of fitness assays conducted between plasmid hosts and a plasmid-free TmR-strain that are otherwise isogenic. The results are shown per plasmid host (x-axis). Data of individual replicates is shown with purple dots (n=5). The mean relative fitness is shown with a black dot and error-bars of x2 standard error of the mean (SEM). The dashed red line marks the expectation for plasmids having a negligible effect on their host fitness (w=1). The mean relative fitness effect of each plasmid variant was compared to 1 using a one-sample, two-sided t-test ($H_0$: w=1; $H_1$: w≠1) with $\alpha$=0.05 using R. The resulting P-value is shown for two plasmid variants where the $H_0$ was rejected (marked also with *).

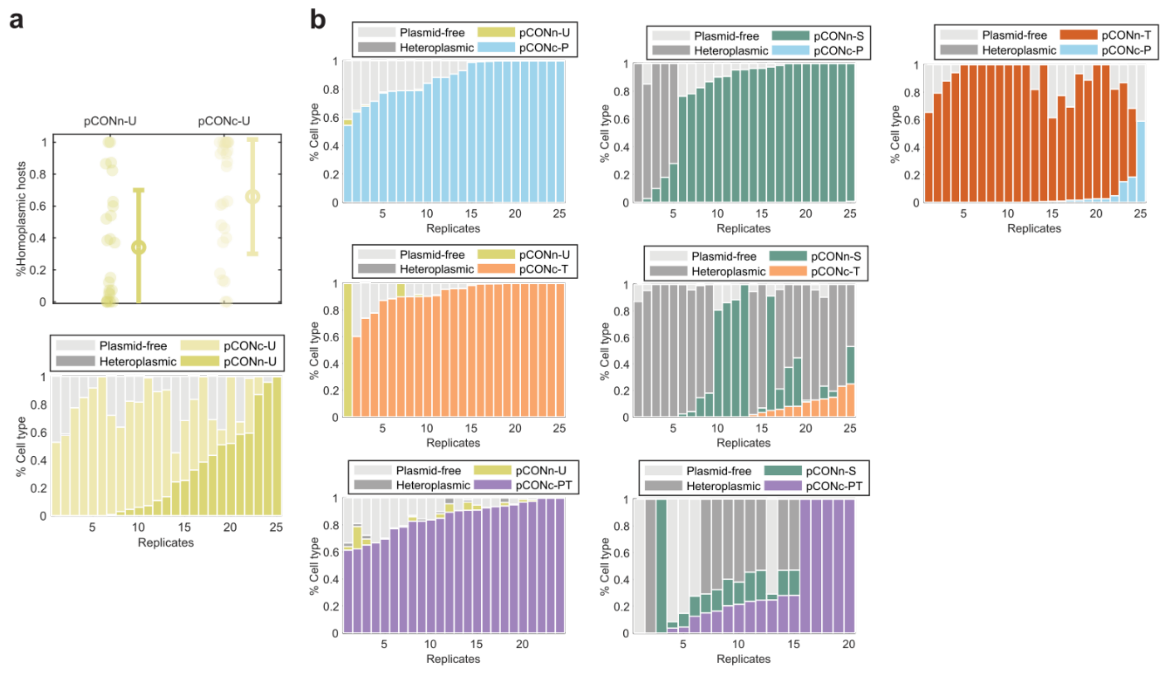

**Fig. S2. Unstable plasmids competitions and results of pairwise competitions with detailed cell type frequencies.**

a) Results of the competition between the two unstable model plasmids pCONn-U and pCONc-U. The competition results presented in the top graph as a scatter plot. Host frequencies are shown with dots. The mean frequency is shown with a circle, with error bars corresponding to standard deviation. The bottom graph shows the distribution of cell types in all competition replicates. b) Cell frequencies in the head-to-head competitions presented in the manuscript Figure 2. Each stacked bar graph shows the results of pairwise plasmid competitions for a different plasmid combination. Bars in the plots correspond to a replicate population with shaded area in the bar proportional to the proportion of cell types in the population at the end of the competition.

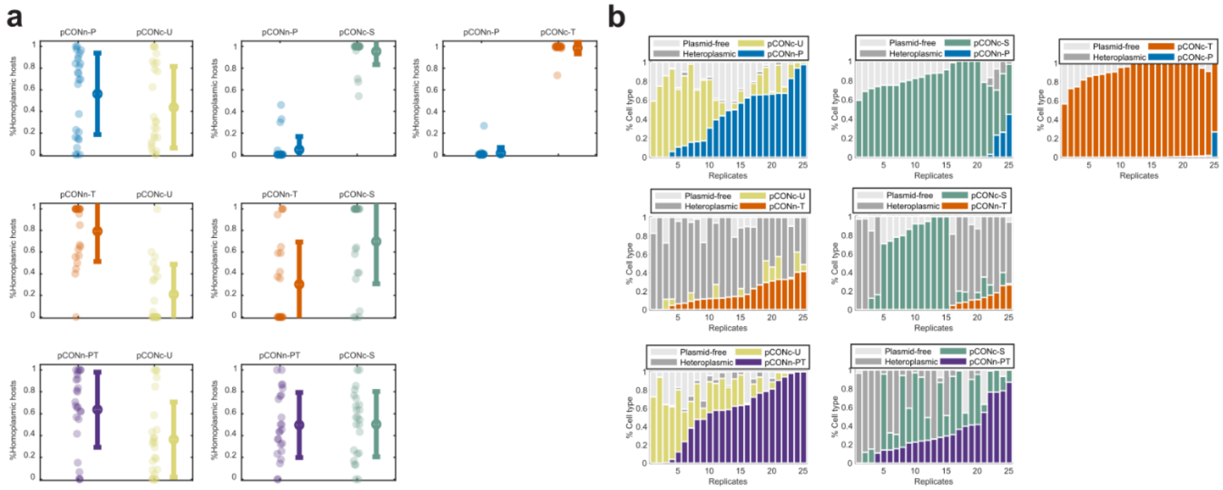


**Fig. S3. Results of reciprocal pairwise competitions.**

Results of the pairwise competitions between model plasmids where the stability systems are exchanged between the plasmid backbones. a) The competition results presented as a scatter plot. Host frequencies are shown with dots. The mean frequency is shown with a circle, with error bars corresponding to standard deviation. b) The competition results presented in stacked bar graphs. Bars in the plots correspond to a replicate population with shaded area in the bar proportional to the proportion of cell types in the population at the end of the competition.

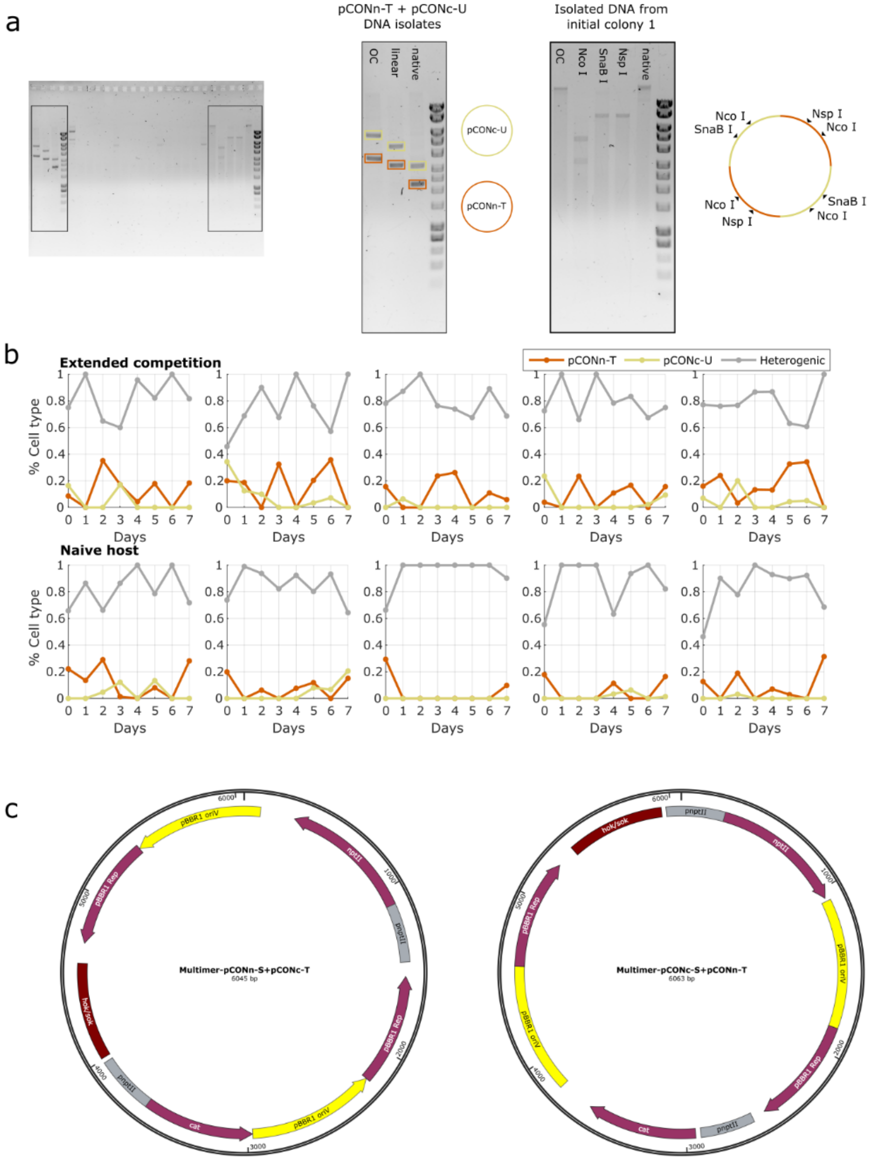


**Fig. S4. Emergence of stable heteromultimers in the competition between T, U and S pCON derivatives.**

The competition between pCONn-T and pCONc-U or pCONc-S yielded a high frequency of heteroplasmic hosts (Fig. S4).

a) To evaluate the plasmid repertoire of heteroplasmic hosts, the plasmid composition within one replicate was analyzed using restriction enzymes. Plasmid DNA was extracted from initial colonies of the competitions between pCONn-T and pCONc-U. The plasmid DNA was nicked with Nb.BsrD I or linearized using Nco I (*cat* and *nptII*), Nsp I (*nptII*) or SnaB I (*cat*). The analysed plasmid DNA revealed the presence of a heteromultimer consisting of two copies of pCONc-U and two copies of pCONn-T.

b) In order to observe the plasmid dynamics over a longer frame of time, either a colony originating from the double plasmid host population of the initial populations or the initial colonies themselves were used to inoculate non-selective LB liquid cultures. After each day the cultures are used to inoculate the next LB liquid cultures with a 1:500 bottleneck. At each day the current liquid culture is diluted in a serial manner and differentially plated in order to obtain the proportions of hosts for the population. The results reveal a stable persistence of plasmid multimers over time (as previously reported for pCON derivatives in Hülter et al. 2020). The presence of multimers was furthermore tested via transformation of the evolved plasmids into a naïve host. Plasmid DNA was extracted from the last timepoint of one replicate of the seven-day assays and used to transform naïve *E. coli recA*::*tetA* cells. The transformed cells were treated according to the transformation protocol described previously. Of the confirmed transformants five colonies were chosen to be observed over the course of seven days. The five replicates were treated as described in the seven-day assay. The results reveal a stable persistence of plasmid multimers over time.

c) The evaluate the presence of heteromultimers in heteroplasmic hosts in the competitions of T and S plasmid derivatives, the plasmid DNA from selected evolved repliactes was sequenced using Oxford Nanopore technology. The sequencing results revealed the presence of heteromultimers consisting of either pCONc-S and pCONn-T or pCONn-S and pCONc-T.

### Supplementary Tables

The tables are supplied in the supplementary data excel file.

**Table S1. Copy number of models plasmids.**

The table supplies the underlying data for the calculation of the plasmid copy number (PCN). The PCN was calculated based on the mean sequencing depth of the plasmid DNA in relation to the mean sequencing depth of the chromosomal DNA. The PCN for the pBBR1 backbone was determined by calculating the average of the model plasmid PCNs.

**Table S2. Oligonucleotides used in this study.**

Names and sequences of the oligonucleotides used as PCR primers in this study.

**Table S3. Full list of identified partitioning systems in the plasmid dataset.**

This table contains a list of partitioning proteins along with associated metadata. Partitioning systems were identified in a plasmid dataset from *Escherichia*, *Salmonella* and *Klebsiella* strains.

**Table S4. Full list of toxin-antitoxin systems for screening.**

This table contains the list of toxin-antitoxin proteins/RNAs from TADB, which were used to screen a plasmid dataset from *Escherichia*, *Salmonella* and *Klebsiella* strains to identify potential TA loci.

**Table S5. Conserved syntenic blocks (CSBs) in X3 plasmids.**

The table contains information on the CSB containing the *agrB-dqlB* locus in PTU-X3 plasmids.

### Supplementary Datasets

The datasets are supplied in the supplementary data excel file.

**Dataset S1. Model plasmid fitness effect on the host.**

The Malthusian parameter was estimated for each plasmid carrying strain in comparison to a trimethoprim resistant strain. The resulting relative fitness of the plasmid hosts is supplied for all replicates.

**Dataset D2. Quantification of plasmid loss.**

Details on the ratio of plasmid free cells and plasmid hosts for each day of the conducted loss assay for all model plasmids.

**Dataset D3. Results of the head-to-head competitions between the pCON-U variants and the stabilized plasmid variants.**

Population size and ratio of cell types present after the competitions between pCON-U variants and the competing plasmid.

**Dataset S4. Results of the head-to-head competitions between the pCON-S variants and the stabilized plasmid variants.**

Population size and ratio of cell type ratios observed after the competitions between the pCON-S variants and the competing plasmids.
